# Neurosecretory protein GL stimulates feeding behavior and fat accumulation in Japanese quails (*Coturnix japonica*)

**DOI:** 10.1101/2025.03.18.644069

**Authors:** Masaki Kato, Eiko Iwakoshi-Ukena, Megumi Furumitsu, Yuki Narimatsu, Kazuyoshi Ukena

## Abstract

Neurosecretory protein GL (NPGL) is a novel hypothalamic neuropeptide that promotes fat accumulation in rats, mice, and neonatal chicks. However, its role in Japanese quails (*Coturnix japonica*) remains unclear. In this study, we investigated the effects of chronic intracerebroventricular infusion of NPGL on body mass, food intake, and fat accumulation in five-week-old male Japanese quails. A 13-day administration of NPGL significantly increased body mass, food intake, and the masses of subcutaneous fat, abdominal fat, and liver. In contrast, water intake and the masses of the pancreas, testes, heart, and muscle remained unchanged. Blood concentrations of triglyceride, glucose, and non-esterified fatty acid were unaffected. Real-time qPCR analysis revealed a significant upregulation of *NPGM*, a paralogous gene of *NPGL*, in the hypothalamus. Additionally, the expression of stearoyl-CoA desaturase 1 (SCD1), a key enzyme in lipogenesis, showed an increasing trend in the liver. Although the fatty acid ratio did not change in the SCD1 activity index (palmitoleate/palmitate), the *de novo* lipogenesis index (palmitate/linoleate) exhibited an upward trend in both the liver and abdominal fat. These results suggest that NPGL promotes fat accumulation in Japanese quails, highlighting its potential role as a contributing factor to obesity in avian species.

## 1. Introduction

Feeding behavior in vertebrates is regulated by bioactive peptides secreted from the central nervous system (CNS) and peripheral tissues (Morton et al., 2006). Among them, many neuropeptides associated with feeding behavior are expressed in the hypothalamus (Valassi et al., 2008). Notably, hypothalamic neuropeptides identified in various vertebrates often exhibit different effects on feeding behavior in birds (Tachibana and Tsutsui, 2016). This discrepancy is particularly evident in orexigenic factors. While orexin (ORX) and melanin-concentrating hormone (MCH) stimulate feeding behavior in mammals, they have no effect in chicks (Ando et al., 2000; Furuse et al., 1999). Similarly, growth hormone-releasing hormone (GHRH) enhances growth and feeding behavior in mammals (Feifel and Vaccarino, 1994) but suppresses food intake in chicks (Furuse et al., 2001). These findings suggest that the regulatory mechanisms of feeding behavior in birds are distinct from those in mammals, and yet unknown factors may play crucial roles in avian species.

In our previous study, we aimed to identify novel regulators involved in feeding behavior and growth in the chicken hypothalamus. We discovered a cDNA encoding a small neurosecretory protein (Ukena et al., 2014). The predicted precursor protein contains a signal peptide sequence, a mature protein sequence of 80 amino acid residues, an amidation signal, and cleavage site. Based on its predicted C-terminal Gly-Leu-NH_2_ sequence, we designated this protein as neurosecretory protein GL (NPGL). Localization studies have shown NPGL-expressing cells across various species, including birds, mammals, and fish. In chickens, *NPGL* mRNA is expressed in the median mammillary nucleus and infundibular nucleus (Ukena et al., 2014). In rodents, *NPGL* mRNA-expressing and immunoreactive cells are localized in the lateroposterior part of the arcuate nucleus (ArcLP) and the ventral tuberomammillary nucleus in rats, but only in the ArcLP in mice (Iwakoshi-Ukena et al., 2017; Matsuura et al., 2017). In fish, *NPGL* mRNA are expressed in the anterior periventricular pretectal nucleus of the anterior preoptic area and lateral tuberal nucleus of the hypothalamus in tilapia (*Oreochromis niloticus*) (Huang et al., 2022). These findings indicate that while NPGL is localized in the hypothalamus across species, the specific neuronal nuclei vary among species. The physiological function of NPGL has been explored through chronic infusion and precursor gene overexpression in the brains of chickens and rodents. Previous studies demonstrated that NPGL promotes food intake and fat accumulation in rats, mice, and juvenile chicks (Iwakoshi-Ukena et al., 2017; Narimatsu et al., 2022; Shikano et al., 2018b, 2019). These findings indicate that NPGL is a novel regulator of energy metabolism in vertebrates.

In chicks, the expression level of NPGL increases with growth, suggesting that the physiological role of NPGL may change over time (Shikano et al., 2018a). However, studies on NPGL in birds have been performed through chronic intracerebroventricular (i.c.v.) infusion in juvenile chicks, leaving the function during puberty and adulthood unexplored. The Japanese quail (*Coturnix japonica*) serves as an excellent model for studying the physiological roles of neuropeptides in birds, particularly during puberty and adulthood, due to its small size and early sexual maturity. Our previous research revealed that the amino acid sequence of mature NPGL in Japanese quails shares 99% homology with that in chickens. Additionally, *NPGL* mRNA-expressing cells in quails are localized in the infundibular nucleus and median eminence (Kato et al., 2024). Moreover, the expression level of *NPGL* mRNA is increased after 24 hours of fasting in male quails (Kato et al., 2024). Despite these insights, the physiological function of NPGL in Japanese quails of puberty remains to be clarified. Therefore, in this study, we investigated the effects of chronic i.c.v. infusion of NPGL in five-week-old male Japanese quails.

## 2. Materials and Methods

### 2.1. Animals

Fertilized eggs of Japanese quail (*Coturnix japonica*) were obtained from a commercial supplier (Motoki Corporation, Saitama, Japan) and incubated in the laboratory. After hatching, male quails were housed in a windowless room at 25°C under long-day conditions (16 h of light and 8 h of darkness) with ad libitum access to water and a commercial diet (Dream Co., Ltd., Aichi, Japan). The experimental procedures were conducted in accordance with the Guide for the Care and Use of Laboratory Animals prepared by Hiroshima University (Higashi-Hiroshima, Japan). The Institutional Animal Care and Use Committee of Hiroshima University approved the protocols (permit number C22-27).

### 2.2. Production of NPGL

Quail NPGL was synthesized using fluorenylmethyloxycarbonyl (Fmoc) chemistry and a peptide synthesizer (Syro Wave; Biotage, Uppsala, Sweden), following our previous methods (Masuda et al., 2015; Shikano et al., 2019). The NPGL sequence was obtained from our prior study (Kato et al., 2024). The purity of the synthesized NPGL was confirmed to be >95% by high-performance liquid chromatography (HPLC). Lyophilized NPGL was weighed using an analytical balance (AP125WD; Shimadzu, Kyoto, Japan).

### 2.3. Chronic i.c.v. infusion of NPGL

The dose of NPGL was set at 15 nmol/day based on previous studies. NPGL was dissolved in 30% propylene glycol. Five-week-old male quails of puberty received i.c.v. infusions of either vehicle (30% propylene glycol) or 15 nmol/day NPGL via an infusion cannula (model 328OP; Plastics One, Roanoke, VA, USA) connected to an Alzet mini-osmotic pump (model 2002, delivery rate 0.5 μl/h; DURECT Corporation, Cupertino, CA). The infusion cannula tip was implanted into the lateral ventricle at the following coordinates: 2.0 mm rostral to the lambda, 1.0 mm lateral to the midline, and 5.5 mm ventral to the skull surface. Osmotic mini-pumps were implanted subcutaneously in the neck. Body mass, food intake, and water intake were measured daily (between 8:00– 10:00) throughout the experiment. After 13 days, the quails have become sexually maturated and were sacrificed by decapitation between 12:00–15:00, and masses of subcutaneous fat, abdominal fat, gizzard fat, liver, pancreas, testes, heart, pectoralis major muscle, pectoralis minor muscle, and biceps femoris muscle were measured. The hypothalamic infundibulum, abdominal fat, and liver were immediately snap-frozen in liquid nitrogen and stored at −80°C for real-time qPCR or gas chromatography-mass spectrometry (GC-MS). Blood samples were collected at the endpoint and centrifuged at 3000×g for 15 min at 4°C to separate the serum.

### 2.4. Serum biochemical analysis

Serum glucose levels were measured using the GLUCOCARD G+ meter (Arkray, Kyoto, Japan). The concentrations of free fatty acids were determined using the non-esterified fatty acid (NEFA) C-Test Wako (Wako Pure Chemical Industries, Osaka, Japan). Serum triglyceride (TG) levels were assessed with the Triglyceride E-Test Wako (Wako Pure Chemical Industries).

### 2.5. Determination of triglyceride concentration in the liver

Lipid extraction from the liver was performed using chloroform:methanol mixture (2:1). TG concentrations in the liver were determined using a colorimetric assay with the Triglyceride E-Test Wako (Wako Pure Chemical Industries).

### 2.6. Fatty acid analysis

Stearoyl-CoA desaturase 1 (SCD1) is an enzyme that converts saturated fatty acids to unsaturated fatty acids. The SCD1 activity index was determined by calculating the palmitoleate-to-palmitate ratio (16:1/16:0) and the oleate-to-stearate ratio (18:1/18:0). The *de novo* lipogenesis (DNL) index was calculated by assessing the palmitic-to-linoleic acid ratio (16:0/18:2n-6). To evaluate endogenous SCD1 activity in liver and abdominal fat, fatty acids were identified using GC-MS analysis (JMS-T100 GCV; JEOL, Tokyo, Japan). Fatty acid extraction was performed according to our previous methods (Narimatsu et al., 2022). The extracted fatty acids were methylated using a Fatty Acid Methylation Kit (Nacalai Tesque, Kyoto, Japan) and purified with a Fatty Acid Methyl Ester Purification Kit (Nacalai Tesque).

### 2.7. Real-time qPCR

RNA was extracted using the RNeasy Micro Kit (QIAGEN, Venlo, Netherlands) for the hypothalamic infundibulum, TRIzol reagent (Life Technologies, Carlsbad, CA) for liver, or QIAzol lysis reagent (QIAGEN) for abdominal fat following the manufacturer’s instructions. First-strand cDNA was synthesized from total RNA (1 μg) using the ReverTra Ace kit (TOYOBO, Osaka, Japan). PCR amplifications were performed using THUNDERBIRD SYBR qPCR Mix (TOYOBO) under the following conditions: 95°C for 20 s, followed by 40 cycles of 95°C for 3 s and 60°C for 30 s using a real-time thermal cycler (CFX Connect; BioRad, Hercules, CA, USA). Primer sets are listed in Table 1. The precursor genes of NPGL, NPGM, proopiomelanocortin (POMC), corticotropin-releasing factor (CRF), neuropeptide Y (NPY), agouti-related peptide (AGRP), and galanin (GAL) were selected as feeding-related factors in the hypothalamus. In the abdominal fat and liver, we analyzed the precursor genes of acetyl-CoA carboxylase (ACC), fatty acid synthase (FAS), SCD1, and peroxisome proliferator-activated receptor γ (PPARγ) as lipogenic factors, along with carnitine palmitoyltransferase 1a (CPT1a), adipose triglyceride lipase (ATGL), and peroxisome proliferator-activated receptor α (PPARα) as lipolytic factors. Insulin-like growth factor 1 (IGF-1) was included due to its role in lipolysis in the liver and its association with adipocyte proliferation or differentiation. Tumor necrosis factor-α (TNF-α), an inflammatory cytokine secreted by chronic obesity was also analyzed.

**Table 1.**
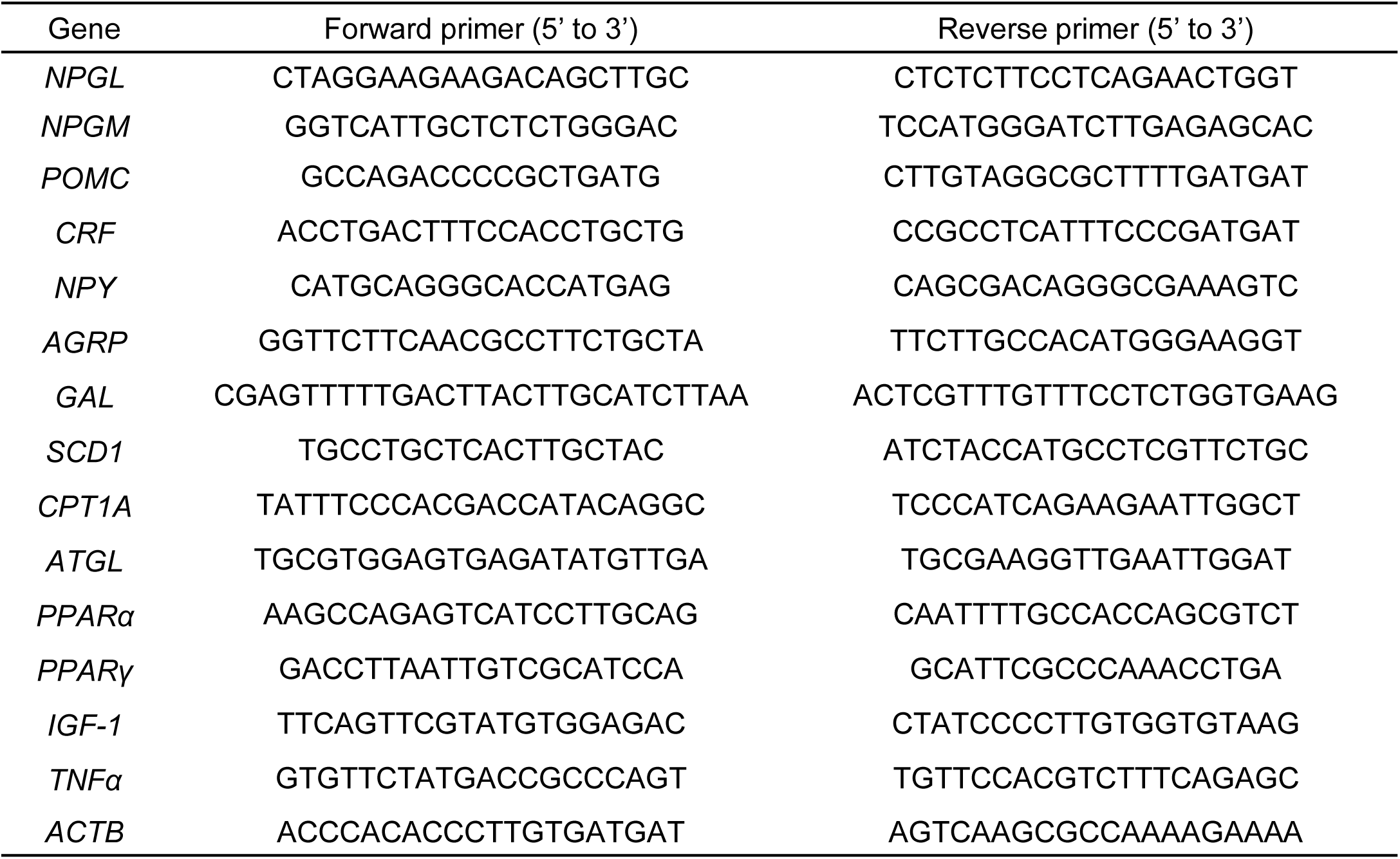
Sequences of oligonucleotide primers for real-time PCR.

### 2.8. Statistical analysis

Differences between the two groups at the endpoint were assessed using the Student’s t-test. Body mass gain and cumulative food and water intake were analyzed using Two-way repeated-measures analysis of variance (ANOVA), followed by Bonferroni’s test. Statistical significance level was set at *P* < 0.05. All results are presented as mean ± SEM.

## 3. Results

### 3.1. Effects of chronic i.c.v. infusion of NPGL on body mass, food intake, and water intake

To evaluate the effects of chronic i.c.v. infusion of NPGL, body mass, cumulative food intake, and cumulative water intake were measured. Chronic i.c.v. infusion of NPGL significantly increased daily body mass gain, resulting in a significant difference in body mass between the two groups at the end of the experiment (Fig. 1A, B). Additionally, cumulative food intake increased significantly from the day before the endpoint (Fig. 1C, D). However, water intake remained unchanged (Fig. 1E, F).

**Figure 1.**
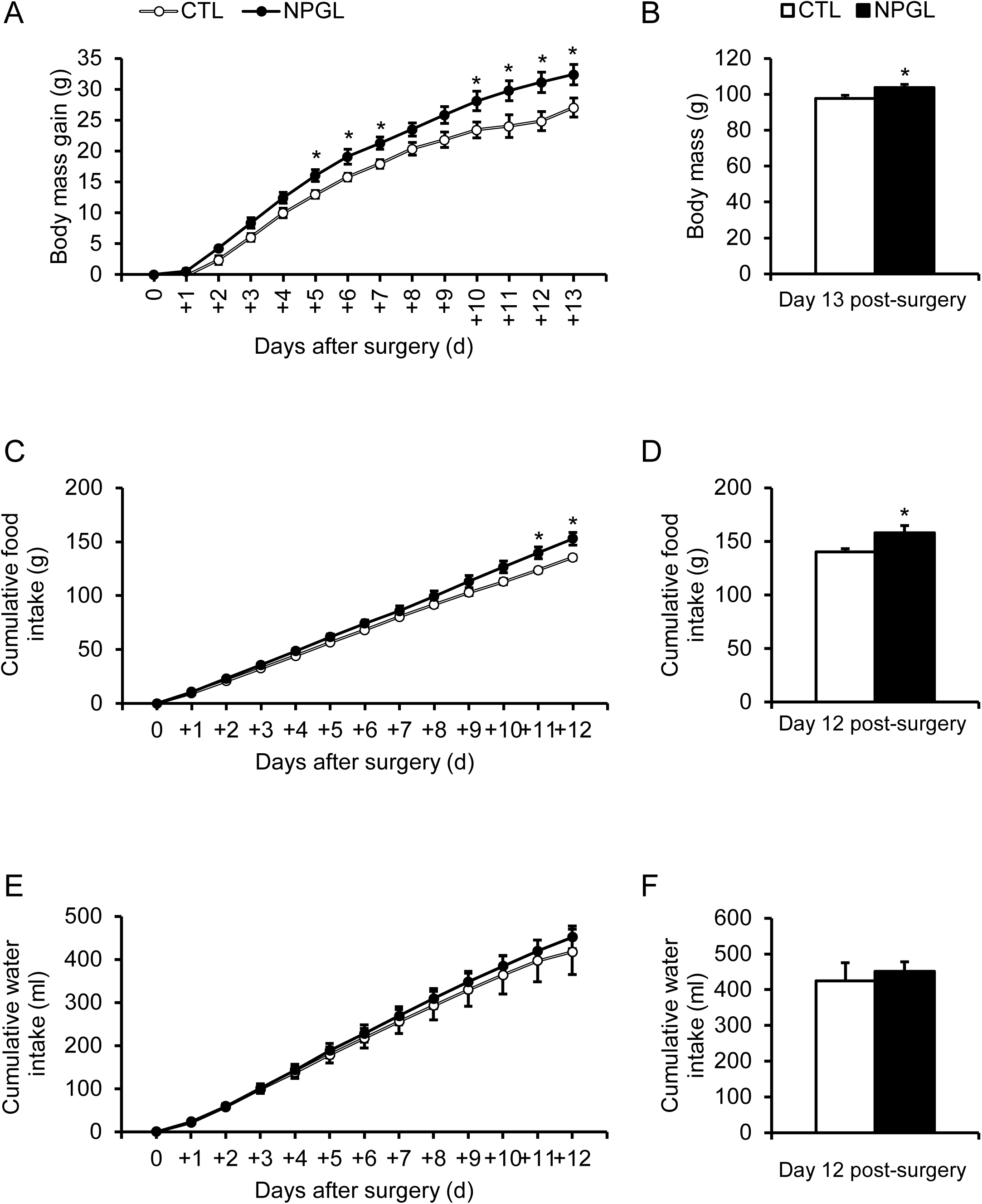
Effect of chronic i.c.v. infusion of NPGL on body mass gain, food intake, and water intake. The results were obtained by the infusion of the vehicle (control; CTL) and NPGL. The change in the body mass gain after surgery (A, B). The cumulative food intake (C, D). The cumulative water intake (E, F). Data are expressed as the mean ± SEM (n = 8–10). Data were analyzed using Student’s t-test and two-way repeated-measures analysis of variance (ANOVA). An asterisk indicates a statistically significant difference (\**P* < 0.05).

### 3.2. Effects of chronic i.c.v. infusion of NPGL on body composition and serum parameters

To assess the effects of chronic i.c.v. infusion of NPGL, tissue masses were measured. Among the adipose tissues, the masses of subcutaneous and abdominal fat were significantly higher in the NPGL-treated group compared to the control group (Fig. 2A). Since fat deposition can also influence the weight of visceral tissues, the masses of the liver, pancreas, testes, and heart were measured. The results showed a significant increase in liver mass (Fig. 2B). In muscles, the masses of pectoralis major and minor muscle were significantly higher in the NPGL-treated group (Fig. 2C). To examine fat accumulation, TG concentrations in the liver and serum, as well as serum glucose and serum NEFA concentrations were measured. However, there were no differences in TG concentrations between the two groups (Fig. 3A, B). Similarly, serum glucose and NEFA concentrations remained unchanged (Fig. 3C, D).

**Figure 2.**
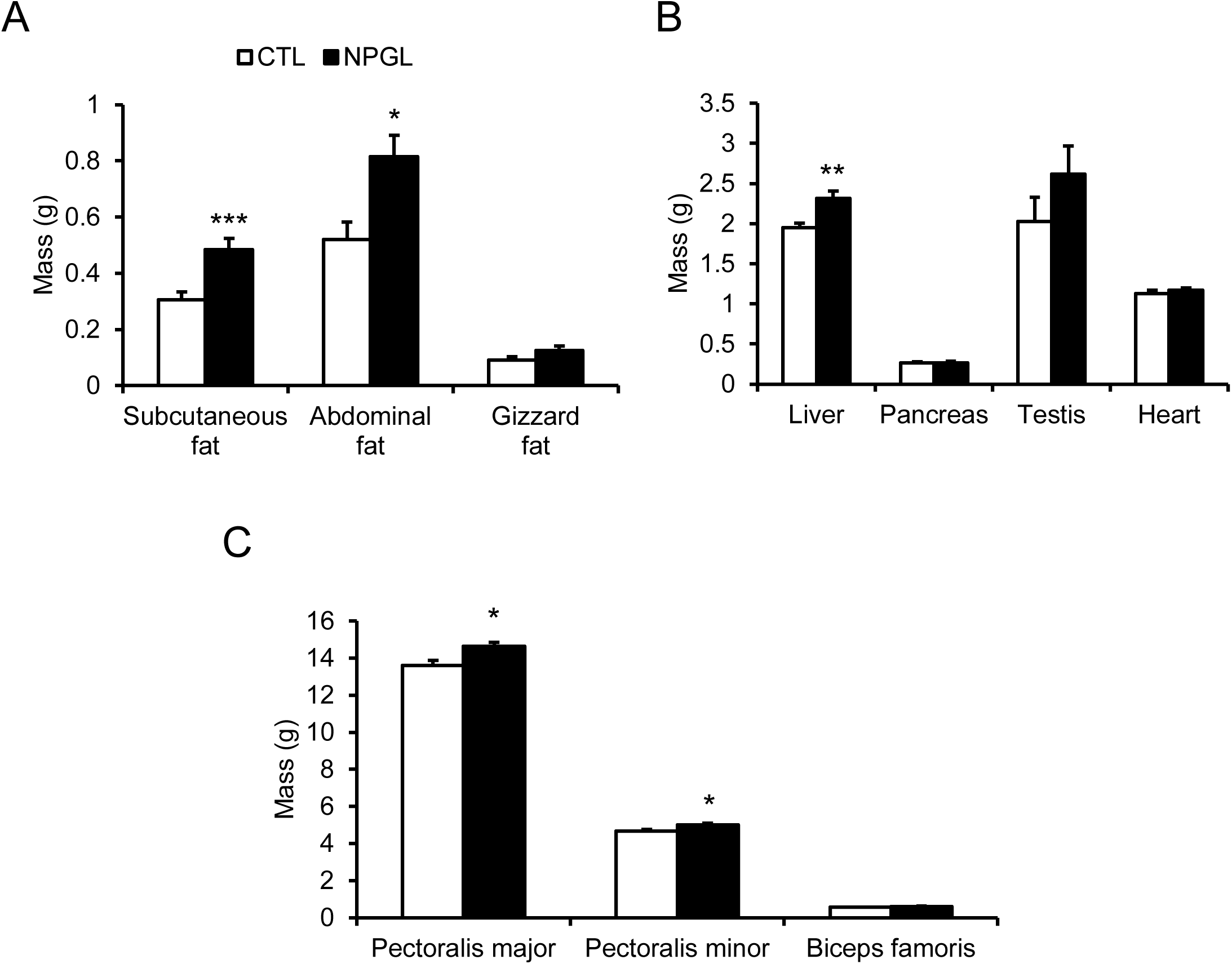
Effect of chronic i.c.v. infusion of NPGL on body composition. The results were obtained by the infusion of the vehicle (control; CTL) and NPGL. The mass of the subcutaneous fat, abdominal fat, and gizzard fat (A). The mass of the liver, pancreas, testis, and heart (B). The mass of the pectoralis major, pectoralis minor, and biceps femoris (C). Data are expressed as the mean ± SEM (n = 8–10). Data were analyzed by Student’s t-test. Asterisks indicate statistically significant differences (\**P*<0.05, \*\**P*<0.01, \*\*\**P* < 0.005).

**Figure 3.**
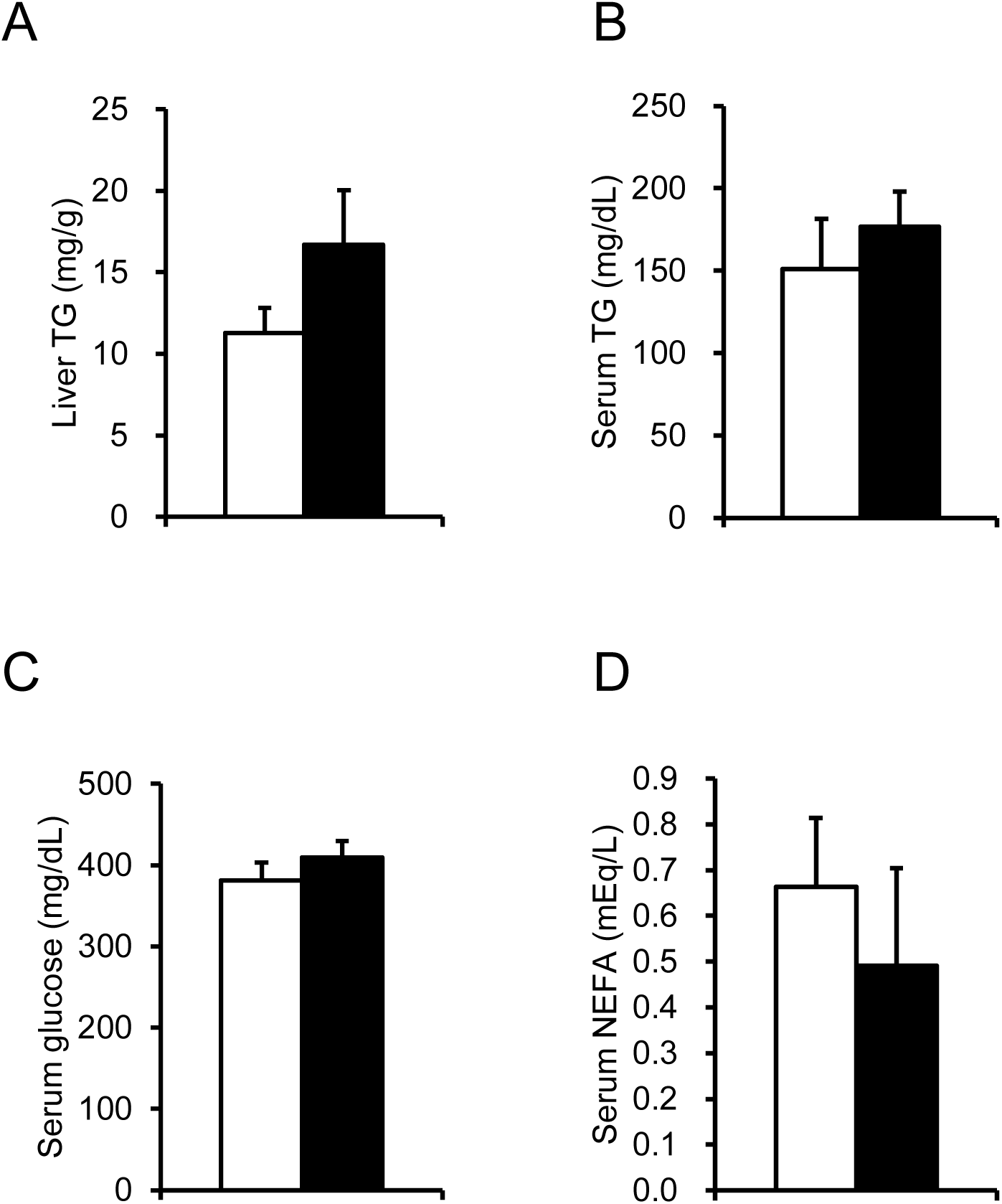
Effect of chronic i.c.v. infusion of NPGL on triglyceride content of the liver and blood component. The results were obtained by the infusion of the vehicle (control; CTL) and NPGL. Triglyceride content of the liver (A). Concentration of triglyceride in serum (B). Concentration of glucose in serum (C). Concentration of non-esterified fatty acid in serum (D). Data are expressed as the mean ± SEM (n = 8–10). Data were analyzed by Student’s t-test.

### 3.3. mRNA expression levels in the hypothalamus

Based on the observed increase in body mass gain and food intake, the mRNA expression levels of feeding behavior-related genes in the hypothalamus were analyzed using real-time qPCR. The results revealed a significant increase in the expression of *NPGM*, a paralogous gene of *NPGL*, while the expression levels of other feeding-related genes, including *POMC*, *CRF*, *NPY*, *AGRP*, and *GAL* remained unchanged (Fig. 4).

**Figure 4.**
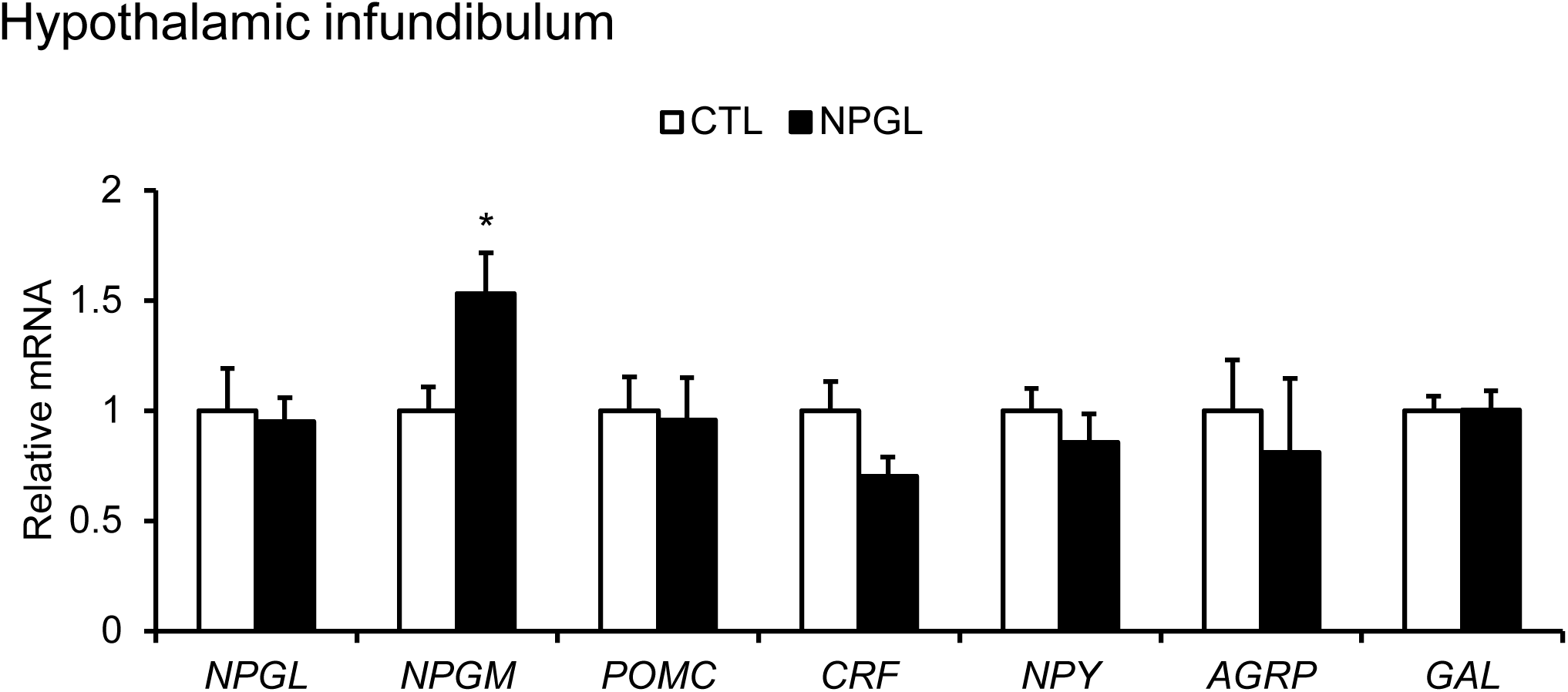
Effect of chronic i.c.v. infusion of NPGL on the mRNA expression of hypothalamic factors [neurosecretory protein GL (NPGL), neurosecretory protein GM (NPGM), pro-opiomelanocortin (POMC), corticotropin-releasing factor (CRF), neuropeptide Y (NPY), agouti-related peptide (AGRP), and galanin (GAL)]. The results were obtained by the infusion of the vehicle (control; CTL) and NPGL. Data are expressed as the mean ± SEM (n = 8). Data are expressed as the mean ± SEM (n = 8–10). Data were analyzed by Student’s t-test. Asterisks indicate statistically significant differences (\**P*<0.05).

### 3.4. mRNA expression levels of lipid metabolic factors in the liver and adipose tissue

To investigate the molecular mechanisms of fat accumulation in the liver and abdominal fat following chronic i.c.v. infusion of NPGL, the mRNA expression levels of lipid metabolic factors were analyzed using real-time qPCR. The mRNA expression levels of *ACC*, *FAS*, *SCD1*, *CPT1A*, *ATGL*, *PPARα*, *PPARγ*, *IGF-1* and *TNF-α* were examined. In the liver, mRNA levels of *SCD1*, which is involved in lipogenesis, showed an increasing trend compared to controls (Fig. 5A). In contrast, no significant changes in gene expression were observed in abdominal fat (Fig. 5B).

**Figure 5.**
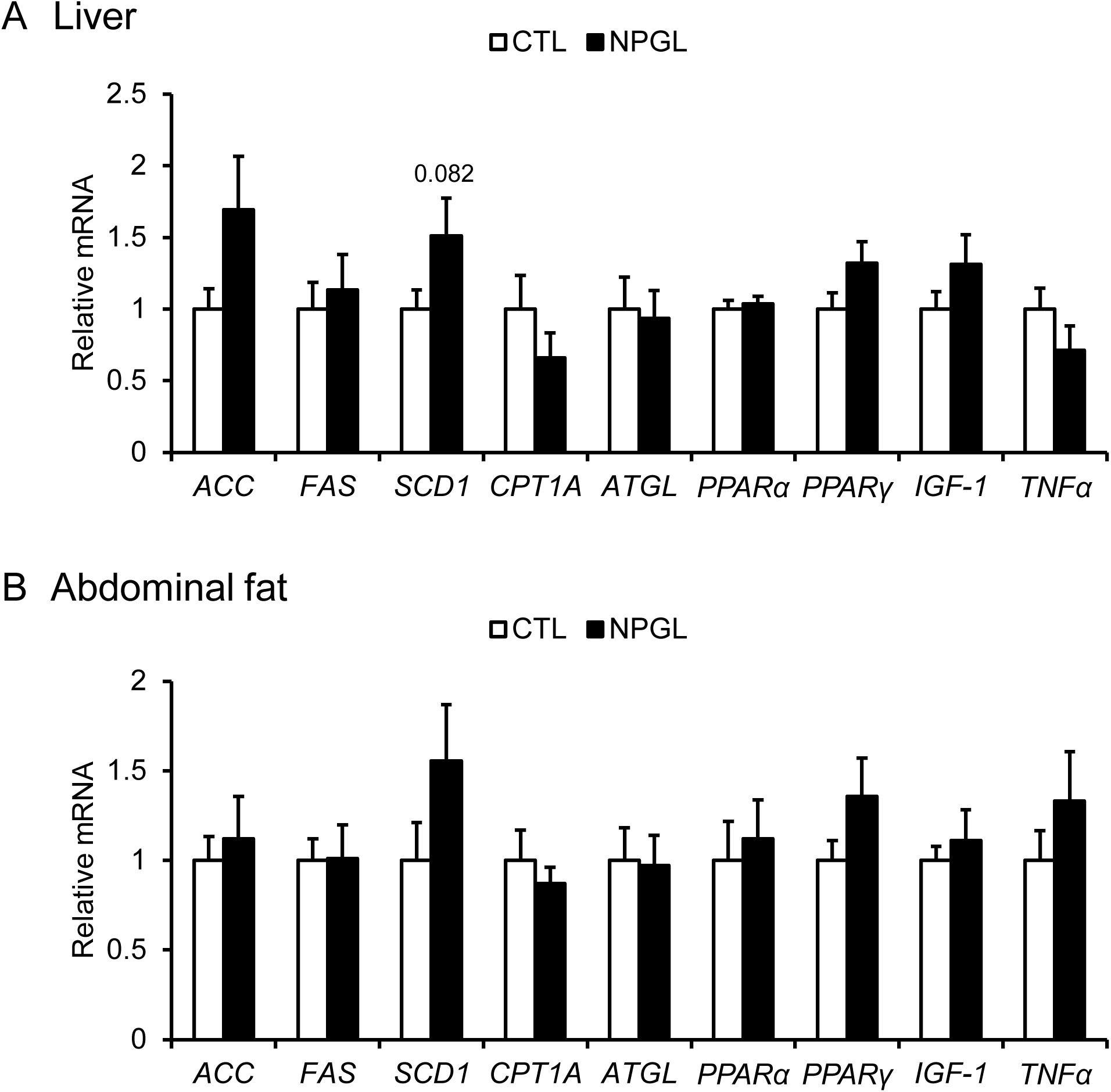
Effect of chronic i.c.v. infusion of NPGL on the mRNA expression of lipogenic and lipolytic factors [acetyl-CoA carboxylase (ACC), fatty acid synthase (FAS), stearoyl-CoA desaturase 1 (SCD1), carnitine palmitoyltransferase 1a (CPT1a), adipose triglyceride lipase (ATGL), peroxisome proliferator-activated receptor α (PPARα), peroxisome proliferator-activated receptor γ (PPARγ), insulin-like growth factor-1 (IGF-1), and tumor necrosis factor α (TNFα)] in the liver and abdominal fat. The results were obtained by the infusion of the vehicle (control; CTL) and NPGL. Data are expressed as the mean ± SEM (n = 8). Data are expressed as the mean ± SEM (n = 8–10). Data were analyzed by Student’s t-test.

### 3.5. Estimation of SCD1 activity index and de novo lipogenesis index by fatty acid analysis

SCD1 is an enzyme that converts saturated fatty acids to monounsaturated fatty acids. To assess the SCD1 activity index, GC-MS analysis was performed on liver and adipose tissues. The palmitoleate-to-palmitate ratio (16:1/16:0) and the oleate-to-stearate ratio (18:1/18:0) showed no significant differences between the two groups (Fig. 6A, B, D, E). However, the DNL index, calculated as the palmitic-to-linoleic acid ratio (16:0/18:2n-6), showed an increasing trend in the NPGL-treated group (Fig. 6C, F).

**Figure 6.**
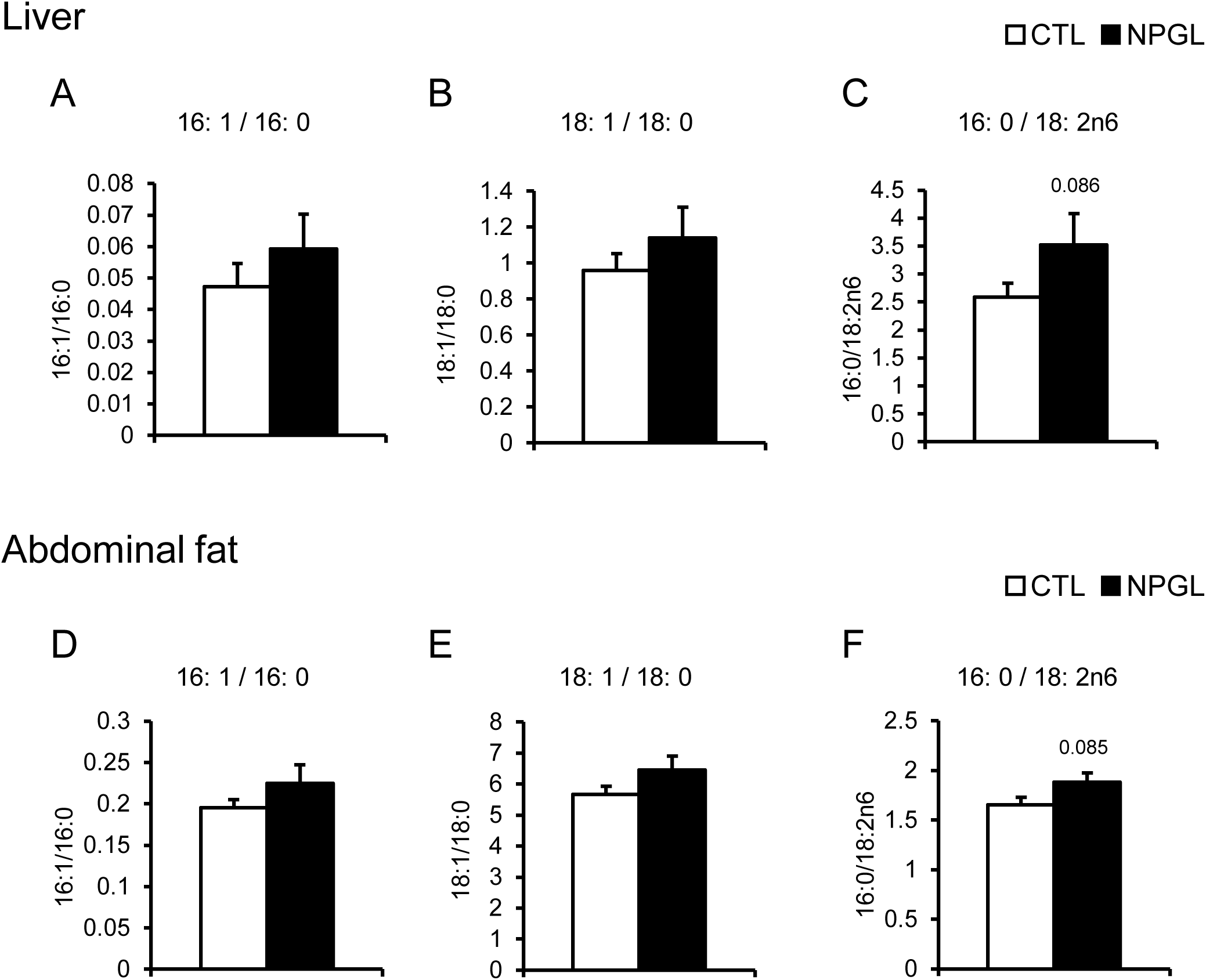
Effect of chronic i.c.v. infusion of NPGL on fatty acid ratio in the liver and abdominal fat. The results were obtained by the infusion of the vehicle (control; CTL) and NPGL. Ratio of fatty acids (16:1/16:0) (A, D). Ratio of fatty acids (18:1/18:0) (B, E). Ratio of fatty acids (16:0/18:2n-6) (C, F). Data are expressed as the mean ± SEM (n = 8–10). Data were analyzed by Student’s t-test.

## 4. Discussion

Physiological analysis of NPGL in birds has previously been limited to juvenile chicks and has not been conducted during puberty and adulthood. In this study, we analyzed the physiological function of NPGL at sexual maturity in quails for the first time. In this study, we examined the phenotypic effects of NPGL on male Japanese quails of puberty by chronic i.c.v. infusion for 13 days. Our data showed that the chronic i.c.v. infusion of NPGL increased body mass and cumulative food intake. In addition, the masses of subcutaneous fat and abdominal fat, liver, and pectoralis major and minor muscles were significantly increased. Furthermore, gene expression analyses and GC-MS analyses indicated a trend toward enhanced DNL in the liver and adipose tissue. The effect of fat accumulation of NPGL has been previously reported in rats, mice and chicks (Fukumura et al., 2021; Iwakoshi-Ukena et al., 2017; Narimatsu et al., 2022; Shikano et al., 2019). In this study, NPGL also induced fat accumulation in Japanese quail, consistent with its effects in other species.

In chickens, disruption of the hypothalamic region has been repotrted to promote overeating, fat accumulation, and atrophy of the testes or ovaries (Jaccoby et al., 1994; Snapir and Robinzon, 1989). Thus, it is speculated that the deficiency of reproductive hormones leads to the hypothalamic-driven overeating and fat accumulation. However, in the present study, testis mass was not affected by NPGL administration, suggesting that reproductive hormones are unlikely to be involved in NPGL-induced fat accumulation in quails. In chickens, growth hormone (GH) secretion from the pituitary gland is known to promote lipogenesis in the liver (Hicks et al., 2017). Previous studies using broiler chicks have also reported that GH promotes the secretion of insulin-like growth factor 1 (IGF-1) in the liver, which stimulates the growth of adipose and muscle tissues (Wang et al., 2015). However, the expression of *IGF-1* mRNA in the liver was not altered by NPGL administration in quails. Thus, it is suggested that NPGL does not activate lipogenesis via GH and IGF-1 secretion.

Cumulative food intake was increased by NPGL administration. However, mRNA expression of feeding-related genes in the hypothalamus did not change in the real-time qPCR analysis except for *NPGM*, which showed increased expression following NPGL administration. *NPGM* is a paralogous gene of *NPGL* and has been shown to induce fat accumulation without increasing food intake in chicks (Kato et al., 2021). In addition, our previous study reported that *NPGM* mRNA is expressed in the mammillary nucleus, and upregulated by fasting (Kato et al., 2024). Although i.c.v. administration of NPGM decreases feeding behavior in chicks, a recent study reported that NPGM increases feeding behavior in mice (Martinez et al., 2023; Narimatsu et al., 2023; Shikano et al., 2018a). Thus, it is suggested that NPGM plays a role in energy metabolism in both birds and mammals. However, the neural projection of NPGL to NPGM is still unknown in Japanese quail. Future studies are needed to elucidate the neural network between NPGL and NPGM.

In this study, muscle mass was increased by NPGL administration. Previous experiments in rats, mice, and chicks have reported that NPGL did not increase muscle mass (Iwakoshi-Ukena et al., 2017; Narimatsu et al., 2022; Shikano et al., 2019). Thus, this is the first report showing that NPGL increases muscle mass in vertebrates. Since the increase in muscle mass may be due to an increase in lipids, muscle cells or both, it is necessary to evaluate the ratio of lipids to protein in muscle tissue. Although chickens and quail are closely related species, they are known to differ in the composition of their thoracic musculature (Matsuda et al., 1983; Nikovits et al., 2001). Muscle is composed of muscle fibers, which are classified into two major types: fast-contracting and slow-contracting fibers. Chicken pectoralis major muscle consists of 99% fast-contracting fibers, whereas quail pectoralis major muscle contains a nearly even mix of fast- and slow-contracting fibers (Matsuda et al., 1983). These findings, together with the results of this study, suggest that NPGL may be involved in the increase of slow muscle fibers in quail pectoral muscle.

Previous studies in mammals have reported that NPGL is a hypothalamic factor that promotes DNL in the liver and adipose tissue (Iwakoshi-Ukena et al., 2017; Shikano et al., 2019). DNL is a metabolic pathway that synthesizes fatty acids from carbohydrates, mainly occurring in the liver and adipose tissue in mammals (Song et al., 2018). Hormones such as insulin and leptin are known to regulate DNL in mammals (Attané et al., 2016; Frederich et al., 1995). Insulin stimulates the expression of glucose transporter 4, promoting glucose uptake into the cells (Abel et al., 2001). Leptin, a hormone secreted from adipose tissue that suppresses feeding, also inhibits DNL by downregulating the expression of sterol regulatory element binding protein-1c (SREBP-1c), a transcription factor involved in fat synthesis (Kobayashi et al., 2020). In birds, however, most DNL occurs in the liver, and hyperglycemia due to insulin resistance is the norm (Hermier, 1997). Moreover, recent findings suggest that leptin in chickens does not function as a hormone as it does in mammals (Friedman-Einat and Seroussi, 2019). Thus, the hormonal regulation of DNL likely differs between birds and mammals. NPGL is speculated to be a novel DNL activator because NPGL administration increases the gene expression of *FAS* and *SCD1*, which are involved in DNL, in chickens (Hermier, 1997; Khatun et al., 2017; Shikano et al., 2019). In the present study, *SCD1* gene expression in the liver also showed an increasing trend, suggesting that DNL is likely activated in quails. SCD1 is an enzyme that converts saturated fatty acids (palmitic acid [16:0] and stearic acid [18:0]) into monounsaturated fatty acids (palmitoleic acid [16:1] and oleic acid [18:1]). The activity of DNL and SCD1 can be assessed by fatty acid ratios (Rosqvist et al., 2019). In this study, the DNL index (palmitic acid [16:0] / oleic acid [18:1]) showed an increasing trend in the liver and adipose tissue. However, the SCD1 activity index (palmitoleic acid [16:1] / palmitic acid [16:0], and oleic acid [18:1] / stearic acid [18:0]) did not change in the liver and adipose tissue. These results suggest that NPGL activates lipogenesis and that the fatty acids synthesized by DNL may be in the process of becoming unsaturated. Furthermore, lipogenesis and lipolysis are regulated in the hypothalamus via the sympathetic nervous system (Buettner et al., 2008; Scherer et al., 2011). Therefore, NPGL may be involved in the sympathetic regulation of lipid metabolism. Future studies are needed to identify target sites and neuronal networks related to NPGL-producing neurons in birds.

In birds, annual changes in day length, called photoperiods, provide clues for predicting seasonal physiological and behavioral changes (Stevenson and Kumar, 2017). Japanese quails are used as model animals for studying seasonal rhythms related to migration and reproduction (Majumdar et al., 2023; Yoshimura et al., 2003). This study is the first to demonstrate that NPGL induces fat accumulation in avian species beyond chickens. However, the biological significance of NPGL remains unclear. We speculate that NPGL-induced fat accumulation is related to seasonal rhythms. Migration, triggered by photoperiod changes, requires large energy reserves, leading to increased food intake and fat storage before migration (Sharma et al., 2023). Therefore, NPGL may be involved in the mechanism of pre-migratory fat accumulation. Future studies should explore the relationship between NPGL and photoperiod. In addition, since chickens and quails are important livestock, NPGL-induced fat accumulation has applications in poultry farming, such as controlling excessive obesity.

## CRediT authorship contribution statement

**Masaki Kato:** Writing – original draft, Writing – review and editing, Visualization, Conceptualization, Methodology, Funding acquisition, Investigation, Project administration. **Eiko Iwakoshi-Ukena:** Investigation, Funding acquisition. **Megumi Furumitsu:** Investigation. **Yuki Narimatsu:** Writing – review and editing, Investigation, Funding acquisition. **Kazuyoshi Ukena:** Writing – review and editing, Conceptualization, Project administration, Supervision, Funding acquisition.

## Acknowledgments

We are grateful to Shogo Moriwaki (Hiroshima University) for the experimental support.

## Data Availability

No data was used for the research described in the article.

